# *Arenaviridae* exoribonuclease presents genomic RNA edition capacity

**DOI:** 10.1101/541698

**Authors:** Elsie Yekwa, Chutima Aphibanthammakit, Xavier Carnec, Caroline Picard, Bruno Canard, Sylvain Baize, François Ferron

## Abstract

The *Arenaviridae* is a large family of viruses causing both acute and persistent infections and causing significant public health concerns in afflicted regions. A “trademark” of infection is the quick and efficient immuno-suppression mediated in part by a 3’-5’ RNA exonuclease domain (ExoN) of the Nucleoprotein (NP). Mopeia virus, the eastern African counterpart of Lassa virus, carries such ExoN domain, but does not suppress the host innate immunity. We have recently reported the crystal structure of the Mopeia virus ExoN domain, which presents a conserved fold and active site. In the present study, we show that the ExoN activity rules out a direct link between ExoN activity and alteration of the host innate immunity. We found that the Arenavirus ExoN, however, is able to excise mis-incorporated bases present at the 3’-end of double stranded RNA. ExoN(-) arenaviruses cultured in cells dampened in innate immunity still replicated in spite of a significant reduction in the viral charge over several passages. The remaining ExoN(-) virus population showed an increased base substitution rate on a narrow nucleotide spectrum, linking the ExoN activity to genome editing. Since, the Arenavirus ExoN belongs to the same nuclease family as that of the nsp14 coronavirus ExoN; which has been recently shown to promote viral RNA synthesis proofreading; we propose that Arenavirus ExoN is involved in a “limited RNA editing” mechanism mainly controlled by structural constraints and a low mutational/fitness ratio.

**Author summary:** Only *Arenaviridae* and *Coronaviridae* encode a 3’-5’ RNA exonuclease domain (ExoN) in their genome. This activity is either used to counteract the innate immunity response during viral infection or to ensure genome stability during replication. Mopeia virus (MOPV), the eastern African counterpart of Lassa virus, carries such ExoN domain, but does not suppress the host innate immunity. We studied MOPV ExoN activity both *in vitro and in cellula* to assess the role of ExoN MOPV and found that the Arenaviral ExoN is fully active on dsRNA, and is able like the one of *Coronaviridae* to excise a mismatched base. We measured genetic stability and found evidence of a limited spectrum of RNA synthesis proofreading mechanism, together with a strongly impacted viral replication. We propose that the Arenaviral ExoN is involved in a functional check of the conserved RNA structures of the viral genome.

## Introduction

*Arenaviridae* is a family of viruses that cause chronic infections of rodents and constitutes a reservoir of human pathogens across the world [1]. Already with a global distribution Lymphocytic choriomeningitis virus (LCMV) is the prototypic member of the family; it is one of the most studied virus and still an underestimated threat to human health [2–6]. In South America, Machupo, Guanarito, Junin, Sabia, and Chapare viruses are responsible for hemorrhagic fever [7] while in Africa Lujo [8] and Lassa viruses (LASV) constitutes a major public health concern [9–13]. Indeed, LASV is responsible for several hundred thousand infections per year alone [14]. It is a common endemic infection in West Africa (Sierra Leone, Guinea, Liberia, Nigeria) responsible for hearing loss, tremors and encephalitis [13,15]. Moreover, this endemic infection frequently spikes a high number of Lassa fever cases associated with significant mortality and high morbidity. The last episode started in February 2018, in the Niger delta region, presents a case fatality rate around 25 % [16]. This new epidemic reinforces the trends observed during the recent epidemics in Nigeria and Benin in January 2016 [17,18], indicating an increase in virulence, an expansion of spreading areas and the number of cases [19]. Humans become infected through contact with infected rodent excreta, tissues, or blood. Person-to-person transmission of Lassa fever can also occur particularly in the hospital environment in the absence of adequate infection control measures [20]. Until now, no licensed vaccine is available, and therapeutic options are limited to early administration of ribavirin. Despite its public health significance, and recent major contributions [21–25], *Arenaviridae* biology is still poorly understood.

*Arenaviridae* are negative-sense single-stranded RNA segmented viruses, with a genome consisting of two segments L (~7.2 kb) and S (~3.4 kb). Each segment has an ambisense coding mechanism, encoding two proteins in opposite orientation, separated by an intergenic region (IGR). The L RNA segment encodes a large protein L (~200 kDa) and a small disordered protein Z (~ 11 kDa) [1]. L is a multi-domain protein including in its N-terminus an endonuclease domain followed by a polymerase domain and in its C-terminus a cap binding like domain (for review [26]). Z, which contains a RING finger motif, is a multifunction protein regulating the life cycle of the virus and during budding assembles to form the matrix [27–29]. The S RNA encodes the precursor of mature virion glycoprotein GP-C (75 kDa); that will give after post-translational cleavage GP-1 (40 to 46 kDa) and GP-2 (35 kDa) [30,31]; and nucleoprotein NP (~ 63 kDa) [25,32]. NP forms a polymer protecting the genomic (and anti-genomic) RNA (RNA_v_) [33]. L and NP together with RNA_v_ form an active ribonucleic complex (RNP) for replication and transcription [34]. In addition to this critical function, NP is involved in clearing off the cytoplasm of double stranded RNAs (_ds_RNA), through its C-terminal exonuclease domain (ExoN) [25,32,35–37]. These _ds_RNAs are markers of viral infection in the cell and are triggering host innate immunity response. Indeed, when _ds_RNA is detected by proteins such as retinoic acid-inducible I (RIG-I) or melanoma differentiation-associated 5 (MDA-5), it initiates a signaling pathway that result in the translocation of interferon (IFN) regulatory factor 3 (IRF-3) to the nucleus [38,39]. Then, IRF-3 activates the expression of IFN-α/β, which initiates the antiviral response in the infected cells and primes neighboring cells for a rapid response to viral invasion. From a modular (sequence and structure) perspective, all NP presents a C-terminal ExoN domain (S1 Fig). The South Eastern African counterpart of Lassa Virus is Mopeia virus (MOPV) [40], a non-pathogenic virus. The NP of these two viruses presents a high sequence identity of about 73%, but contrary to Lassa virus, MOPV infection does not result in innate immunity suppression [41,42] leading us to suspect that the domain was not fully functional against _ds_RNA. Recent studies reported the structure of Mopeia ExoN domain [43] and evidence of an ExoN activity in NP of MOPV, essential for multiplication in antigen-presenting cells [44]. The observed fold conservation and activity raises questions about the biological role of the NP-exo MOPV, and whether it could be conserved for other functional or structural reasons [26]. In the DNA world, ExoNs are mainly involved in mechanisms of genome stability and error correction during or after DNA synthesis. Yet, in viral RNA world, the existence of ExoN is of rare occurrence as only two families of viruses possess a 3’-5’ ExoN member of the DEDD super family: *Arenaviridae* and *Coronaviridae* [25,45–47]. The Coronavirus ExoN is part of the nsp14 protein, associated to the main replicative RNA-dependent polymerase nsp12. During RNA synthesis, nsp14 belongs to the replication/transcription complex (RTC), excises mismatched bases occurring during processive RNA synthesis, and contributes to overall RNA synthesis fidelity [47–51].

Having noted the structural and functional relatedness of *Arenaviridae* ExoN to the *Coronaviridae* ExoN, we engaged into mechanistic studies of the MOPV ExoN. Here, we present a detailed characterization of the activity of NP-exo MOPV including substrate specificity and ion dependency, compared to the one of LCMV. We show that the *Arenaviridae* ExoN is active on 3’ mismatched _ds_RNA substrate mimicking a stalled RNA synthesis intermediate, in a remarkable substrate requirement similarity to Coronavirus nsp14. We report, however, that a mutated NP-exo MOPV abrogating the ExoN activity does not lead to an overall higher mutation rate in the surviving viruses, but rather drastically reduces the number of infectious viruses while increasing the release of non-infectious material from infected cells. Interestingly, few nucleotide substitution types appear to be significantly increased in the ExoN(-), establishing that ExoN is active on its own genomic RNA. All together these results confer potentially significant roles to the ExoN domain in the *Arenaviridae* life cycle.

## Material and Methods

### Cloning, mutagenesis, protein production and purification

cDNA corresponding to NP ExoN domain of: MOPV (residues 365-570 - P19239), LCMV (residues 357-559 - NP_694852) were cloned into the pETG20A expression vector using the Gateway® method (Invitrogen), which adds a cleavable thioredoxin-hexahistidine tag at the N-terminus. The integrity of the DNA construct was verified by DNA sequencing. The sequences of the primers used to sub cloned each domain were:

LCMV forward:

GGGGACAAGTTTGTACAAAAAAGCAGGCTTAGAAAACCTGTACTTCCAGGGTTTAAGCTACAGCCAGACAATGCTTTTAAA,

LCMV reverse:

GGGGACCACTTTGTACAAGAAAGCTGGGTCTTATTATGTCACATCATTTGGGCCTCTA,

MOPV forward:

GGGGACAAGTTTGTACAAAAAAGCAGGCTTAGAAAACCTGTACTTCCAGGGTTTAACCTACTCTCAGACAATGGA,

MOPV reverse:

GGGGACCACTTTGTACAAGAAAGCTGGGTCTTATTACAGGACAACTCTGGGA.

Plasmids were used to transformed *E.coli* strain C2566 (NEB) protease-deficient and carrying pRARE plasmid (Novagen). Bacteria were grown in LB medium (AthenaES) at 37°C to an OD600_nm_ of 0.5. Expression was induced with 0.5 mM IPTG, and bacteria were grown shaking at 210 rpm overnight at 17°C in presence of 100µM of ZnCl_2_. Bacteria were pelleted, frozen, and stored at −80°C.

The three domains were purified at 4°C. Frozen pellet were melted on ice, resuspended in lysis buffer (20mM HEPES pH7.5, 300 NaCl, 5 mM imidazole, 5% glycerol, 0,1 mg/ml lysozyme and 50 µg Dnase), sonicated, and the lysate was cleared by centrifugation at 20,000 rpm for 30 min. Each protein was first purified by metal affinity chromatography using 2ml of His pur^TM^ cobalt column (Thermo Scientific). The tag was removed by cleavage with TEV protease followed by purification on a second cobalt affinity chromatography. Proteins were further purified by gel filtration using superdex 75 column (GE Healthcare) in 20 mM HEPES pH 7.5, 300 mM NaCl, 2 mM MgCl_2_ and 5% glycerol.

Mutants were generated for each domain by introducing single point mutations using the Quick change site-directed mutagenesis kit (Stratagene). The primer sequences used for mutagenesis are listed in the supplementary S1 Table. The presence of *ad hoc* mutations and the integrity of the complete coding region of each mutant were confirmed by sequencing. All the mutants were expressed and purified following the established protocol.

### RNA labeling and preparation

Synthetic RNAs used in this study were purchased from Dharmacon or Biomers (HPLC grade). They are listed in S2 Table and their predicted structures are shown in S2 Fig. All sens RNA strands were labeled at their 5’ end with [γ-^32^P] ATP using protein nucleotide kinase (NEB) according to the manufacturer’s instructions. For experiments involving overhang mismatched _ds_RNA, the _ds_RNAs were generated by annealing an anti-sense RNA strand containing 3’-phosphate modifications with its 5’-radioactively labeled sens RNA strand. The annealing condition was, heating at 70 °C for 10 min and then cooling down to room temperature (with a primer/template ratio of 1.2:1).

### Exonuclease activity assay

Reactions were carried out in a buffer containing 20 mM Tris-Hcl, 5 mM MnCl_2_ (unless specified by 5 mM of: MgCl_2_, CaCl_2_, or ZnCl_2_) and 5 mM DTT. Standard reactions contained 0,25 µM of protein (NP-exo MOPV or LCMV or mutants) and 1,25 µM of radiolabeled RNA substrate. After incubated at 37°C, the reactions were quenched at intervals between 0 and 30 minutes by the addition of an equal volume of loading buffer (formamide containing 10mM EDTA). The products were heated at 70°C for 5 minutes, rapidly cooled on ice for 3 minutes followed by separation in a 20% poly-acrylamide gel containing 8 M urea and buffered with 0,5X Tris-borate-EDTA. Gels were exposed overnight to a phosphor screen and then visualized with a phosphoimager FLA-3000 (Fuji). Total RNA degradation products were quantified using Image Guage (Fuji), the speed of cleavage determined and graphs plotted using GraphPad PRISM version 6.0. Experiments were carried out at least in triplicate and only representative gels are shown.

### Thermal shift assay

A real-time PCR set-up (Bio-rad) was used to monitor the thermal unfolding of the ExoN domain of NP-MOPV or -LCMV alone or in the presence of different divalent ions Mn^2+^, Mg^2+^, Ca^2+^ and Zn^2+^. Proteins were equilibrated in a buffer containing 20 mM HEPES pH 7.5, 300 mM NaCl, 5% glycerol. All reactions were set up in a final volume of 25 µl in a 96-well plate with total protein concentration of 1,8 mg/ml, 1x SYPRO Orange and incubated with or without 5 mM of metal ions. The PCR plates were sealed with optical sealing tape (Bio-rad) and incubated in the PCR machine for 2 minutes at 20°C followed by 0,2°C increments to a final temperature of 95°C. Thermal denaturation was monitored using SYPRO Orange (Life Technologies) and the fluorescent intensities were measured at 490 nm excitation and 530 nm emission wavelengths. The unfolding of proteins was monitored by following the increase in fluorescence of the probe as it binds to exposed hydrophobic regions of the denatured protein. The Tm was then calculated as the mid-log of the transition phase of the florescence curve using the Boltzmann equation. All measurements were performed in triplicates.

### Structure and sequence analysis

#### Structure and sequence comparison of Arenavirus exonuclease with other viral exonuclease

Structure similarities were search with PDBeFold [52] using the MOPV exonuclease domain as a search model (PDB:5LRP). Corresponding sequences were aligned based on structure comparison using Expresso [53]. Figures were generated with the programs ESPript-ENDscript [54], WebLogo server [55] and UCSF chimera [56].

#### Sequence retrieval of mammarenavirus L and analysis

All annotated complete protein sequences of Mammarenaviruses L protein were downloaded from NCBI. The dataset of 559 sequences was manually curated using Jalview [57] in order to remove Identical, mis-annotated or complete sequences with undefined amino acid (X). The remaining 395 sequences were aligned using MUSCLE [58] constituting the *Mammarenavirus* (MAMV) dataset. From this latter two dataset are being derived the Mopeia virus dataset of 12 sequences, and the Lassa Virus dataset of 277 sequences, as being the closest homologue of Mopeia virus with a large number of sequences. Amino acid composition (%) for position in the sequence corresponding to the Mopeia emerging mutant were calculated with Jalview and represented using WebLogo [55] for the three datasets.

### Viral infection and genome sequencing

The MOPV strain AN21366 (JN561684 and JN561685) was used to establish the reverse genetics system for MOPV. The detailed procedures for virus rescue, production, titration and infection of Vero E6 cells (VERO C1008 [Vero 76, clone E6, Vero E6] ATCC® CRL-1586™ (ATCC, LGC Standards, Molsheim, France) are described in [44]. In brief, the recombinant NP-exo WT and mutant (D390A/G393A) MOPV were used to infect Vero E6 cells using a MOI of 0.01. Supernatants were collected four days post infection. Viral production/titration/infection were repeated iteratively for 10 times. For viral titration, the presence of the viruses was revealed by immunostaining in infected cells with a polyclonal rabbit antibody that recognizes the MOPV Z protein (Agrobio, France) and a phosphatase alkaline-conjugated polyclonal goat anti-rabbit antibody (Sigma) and 1-Step NBT/BCIP substrate (Thermo Fisher scientific, Waltham, MA). Results are expressed in Focus Forming Unit per mL (FFU/mL). For RNA quantification, viral RNA were extracted from cell culture supernatants (Qiagen, Courtaboeuf, France) and quantification was performed with the EuroBioGreen qPCR Mix Lo-ROX (Eurobio, Les Ulis, France), using an in house developed assay with 5’-CTTTCCCCTGGCGTGTCA-3’ and 5’-GAATTTTGAAGGCTGCCTTGA-3’ primers. Deep sequencing analysis of viral genomes was performed as described in [44].

## Results

### The MOPV NP-exo exhibits a metal-dependent 3’-5’ ExoN activity

The ExoN domain of NP-MOPV (NP-exo MOPV) was incubated with a 5’ radiolabeled 22 nt RNA hairpin (HP4, S2A Fig) whose 3’-end is base-paired into a double stranded RNA. The reaction was stopped at intervals of 0.1, 5 and 30 minutes (Fig 1A). In the presence of Mg^2+^, NP-exo MOPV is able to cleave this stable RNA hairpin (DG=-14.7 kcal/mole) predominantly down to a 18-mer product. After removing the 1^st^ 4 nucleotides, degradation stops at the loop region. A similar experiment in the presence of Mn^2+^ allows further degradation into the loop, whereas ExoN is inactive in the presence of either Zn^2+^ or Ca^2+^. The laddering degradation pattern visualized on the gel, together with radio-label quantification indicate that it acts in the 3’-5’ direction (S2B Fig). Visual examination of degradation kinetics shows, a band-product accumulation prior to G nucleotides, indicating that the latter are slower to remove than Cs. We also tested the ExoN activity of the ExoN domain of NP-LCMV (NP-exo LCMV) (Fig 1A) on the same RNA substrate. Both proteins cleave the RNA following a similar degradation pattern and comparable kinetics (S2B Fig and S3 Table). The two ExoNs exhibit their highest activity in the presence of Mn^2+^, followed by Mg^2+^ and they are inactive in the presence of either Ca^2+^ or Zn^2+^ or EDTA (Fig 1A). These results show that the nature of the ions is altering the ExoN activity and its associated pattern of degradation can be greatly modulated by the nature of the metal ion co-factor.

**Fig 1.**
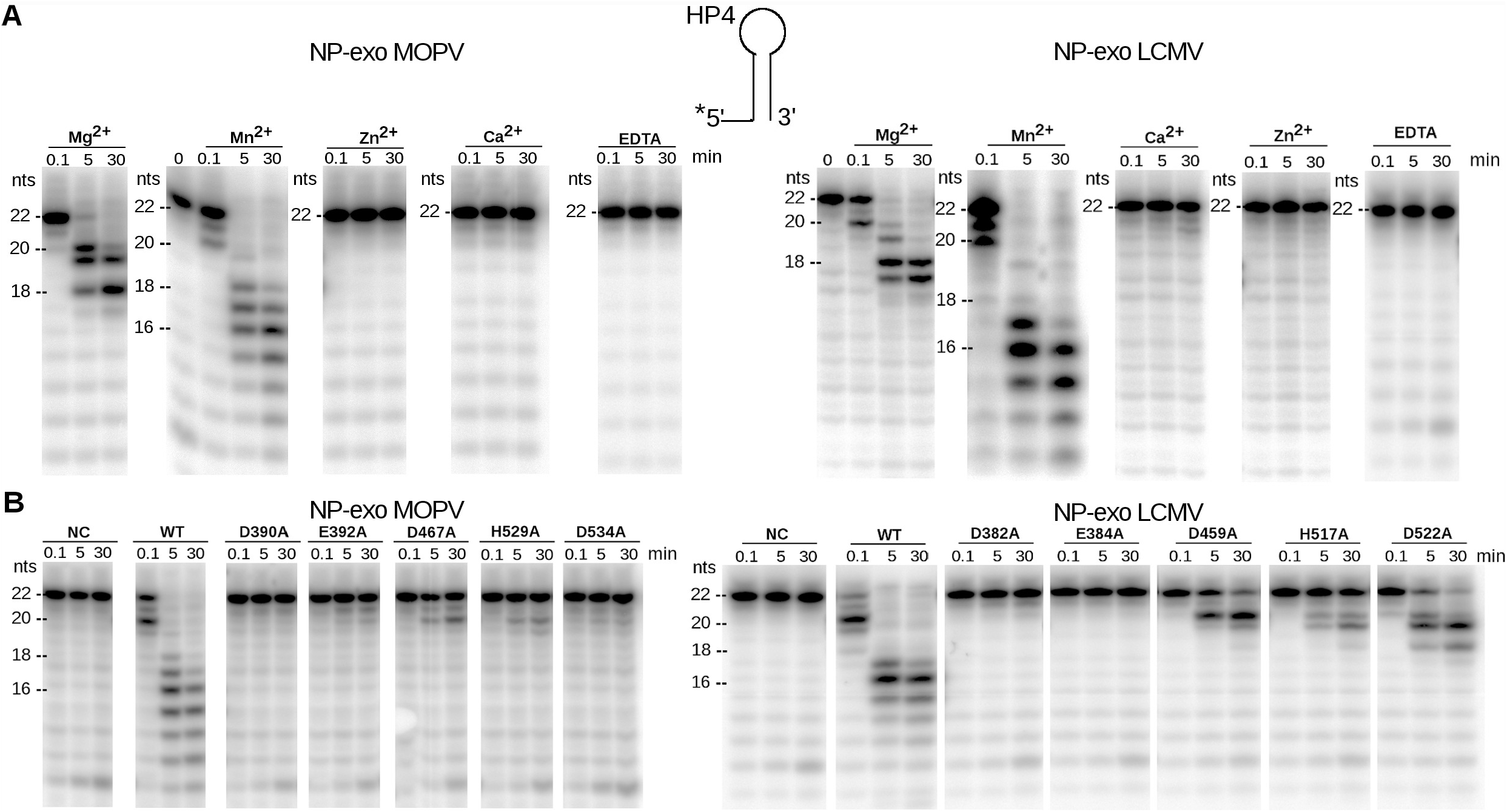
Comparison of the ExoN activity of NP-exo MOPV and NP-exo LCMV. (A) Effect of divalent cations on ExoN activity of NP-exo MOPV and NP-exo LCMV. RNA HP4 was incubated with NP-exo MOPV or NP-exo LCMV for 0, 0.1, 5 and 30 minutes (min) in the presence of 5 mM Mg^2+^, Mn^2+^, Zn^2+^, Ca^2+^, or EDTA. Digestion products were separated on a 20 % denaturing PAGE and revealed by autoradiography. (B) Comparative mutational analysis of ExoN activity. The DEDDh residues were mutated to alanine. Equal amounts of wild type (WT) or mutants of NP-exo MOPV or NP-exo LCMV were incubated with HP4 for 0, 5 and 30 mins. Products were separated on denaturing PAGE and visualized by autoradiography. NC indicates the substrate without proteins. Sketch on the top of figure A illustrates the hairpin structure of HP4. On the side of each gel is presented the migration size ladder in nucleotides (nts).

### Stabilizing effect by ion cofactor is not correlatable to NP-exo activity

To study the effect of divalent metal binding on the stability of NP-exo MOPV, we measured the change in melting temperature (Tm) by a Thermal Shift Assay (TSA) in the presence of 5 mM of several metal ions. NP-exo MOPV without metal ions has a Tm of 49.4 °C. Positive Tm shifts are observed in the presence of MnCl_2_ (+16.5 °C), MgCl_2_ (+4.9 °C), CaCl_2_ (+7.5 °C) and a slight negative shift in ZnCl_2_ (−2.2 °C) (Fig 2). Simultaneously we compared the effect of these ions on the stability of NP-exo LCMV (Fig 2). The Tm value for NP-exo LCMV without metal ions is 50.6 °C and increases in the presence of MnCl_2_ (+13.6°C), MgCl_2_ (+6 °C), CaCl_2_ (+7.67 °C) decreases with ZnCl_2_ (−10.3 °C). These results indicate that the ion stabilization pattern for each ExoN is unique nevertheless, a similar stabilization trend is observed with MnCl_2_ inducing highest stability in all ExoNs. It is also worth noting that CaCl_2_ which inhibits the 3’-5’ ExoN activity is a better stabilizer than MgCl_2_.

**Fig 2.**
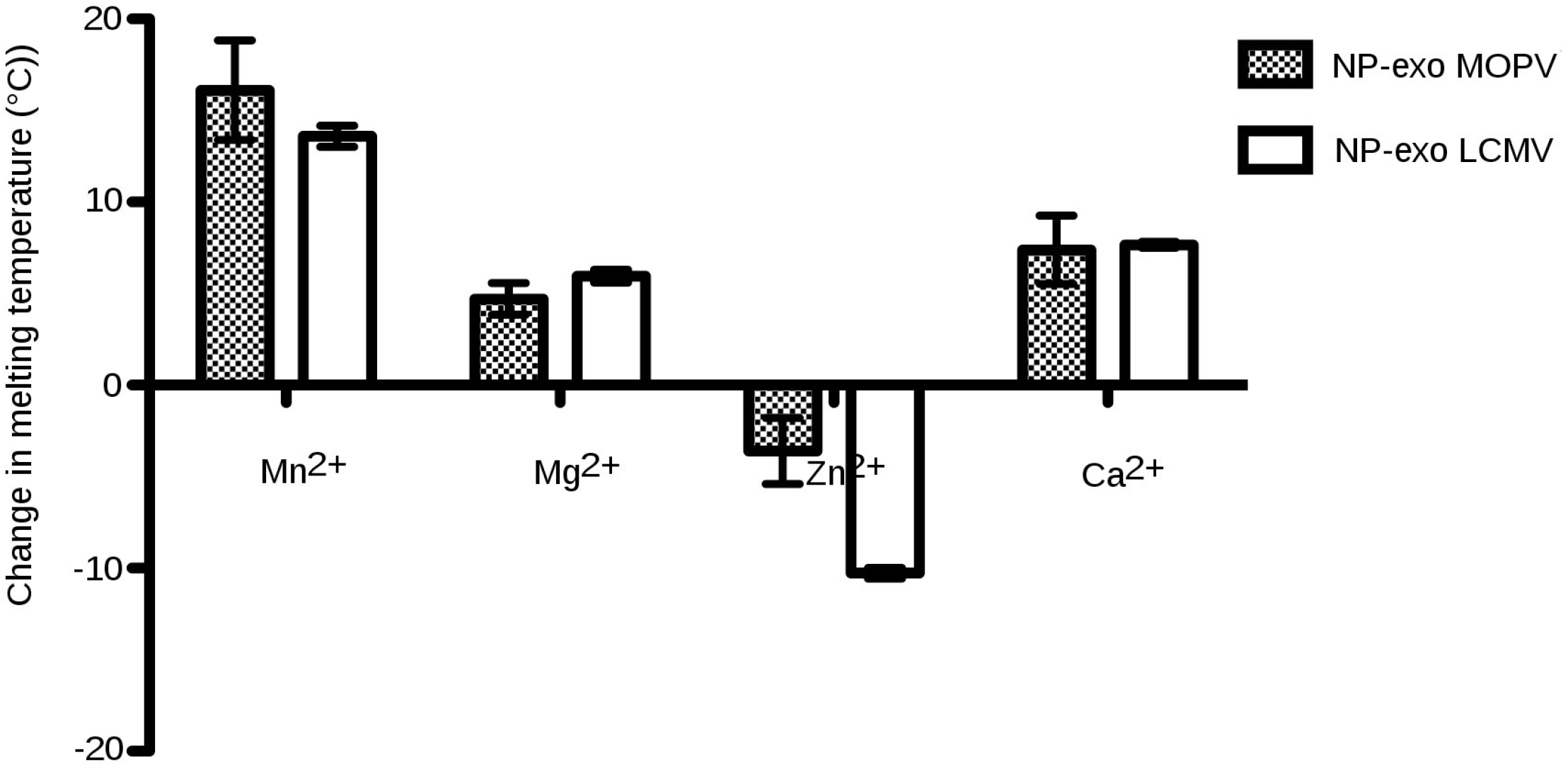
Effect of divalent-Cation on thermal stability of NP-exo MOPV and NP-exo LCMV. Bar chart showing the shifts in melting temperatures of NP-exo MOPV and NP-exo LCMV measured in the presence of 5 mM of the indicated ions by TSA.

Our results show that stability and activity are uncoupled: lowering the energy of the domain is not key for activation. Rather the nature of the ion plays a key role: the small radius and higher coordination of Mn^2+^ over Mg^2+^ allow a higher number of water molecules available for being activated for the nucleophilic attack. On the contrary, Ca^2+^ with a larger radius slightly deforms the catalytic site [43] and increases the distances with the substrate, thus impairing the reaction.

### NP-exo MOPV catalytic residues compared to NP-exo LCMV

We mutated each catalytic residue to alanine in order to assess their respective contribution in the conserved DEDDh catalytic motif, and tested them for ExoN activity. For the NP-exo MOPV mutants D390A, E392A and D534A, the 3’-5’ ExoN activity is completely abolished whereas D467A and H529A are still able to slowly excise up to two nucleotides. For the NP-exo LCMV mutants, a slightly different result is observed. D382A and E384A show a complete loss of activity, D459A excises two nucleotides but more efficiently than D467A of NP-exo MOPV as judged by the diminution of the 22 nts band-product. The H517A also shows residual activity while the D522A is able to degrade almost the total amount of 22 nt dsRNA up to 20/19 nts (Fig 1A). We compared the efficacy of cleavage between the wild type NP-exo MOPV and NP-exo LCMV to their corresponding mutants D467A and D459A respectively. Our kinetic experiment indicates that the initial excision rate of NP-exo MOPV and NP-exo LCMV wild types are similar, the rate of D459A of LCMV decreases to about half that of the wild-type, and that of MOPV D467A is significantly affected (S3 Fig).

### NP-exo MOPV _ds_RNA substrate specificity

In order to investigate the substrate requirement for NP-exo MOPV, NP-exo LCMV, we tested their activities on different RNA substrates HP4, A30 (poly A) and LE19. All these single stranded RNA (_ss_RNA) forms several types of secondary structures RNA, which were predicted using Mfold server [59] (S2A Fig). The ExoN assay confirms and extends findings shown in Fig 1, *i.e* that NP-exo MOPV and NP-exo LCMV cleave RNA substrates whose 3’ ends are engaged into a double stranded structure (Fig 3), consistent with a strict specific double stranded RNA requirement. It is particularly striking in the case of LE19: at time 0, we observed the 3 species of secondary structures (migration for type A:19, B: 18 and C:17 nucleotides respectively) and with time the top band-product disappears to the profit of an RNA of 17 nucleotides. We observed that NP-exo seems to be partly active on small secondary structure _ds_RNA but is inactive on _ss_RNA (Fig 3).

**Fig 3.**
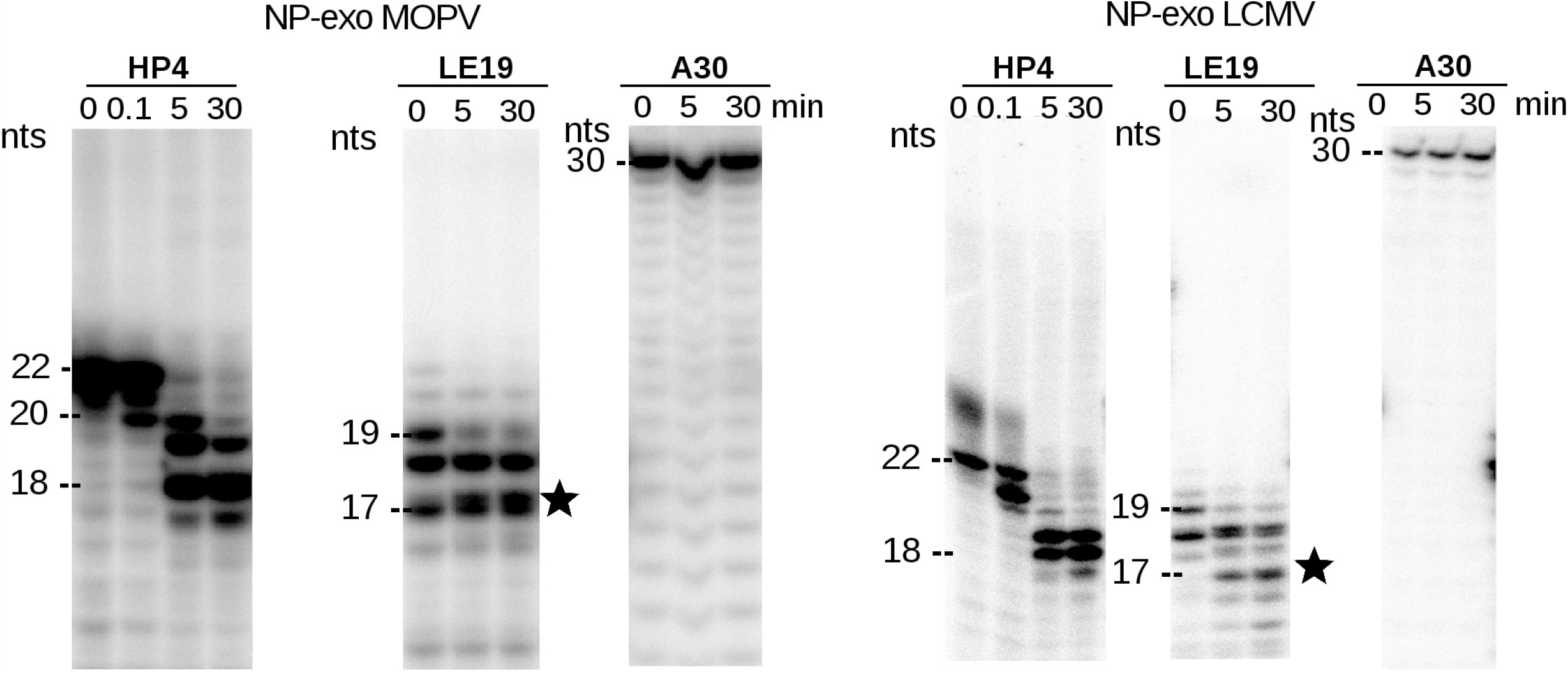
Comparison of the substrate on the ExoN activity of NP-exo MOPV and NP-exo LCMV. Comparative ExoN activity on three different RNA substrate; A30 (ssRNA), LE19 (ssRNA forming three types of secondary structures), HP4 (stable RNA hairpin). Equal amounts of each RNA substrate were incubated with NP-exo MOPV or NP-exo LCMV for 0, 5, 30 minutes. Digestion products were analyzed as in Fig1. On the side of each gel is presented the migration size ladder in nucleotides (nts). A star highlight the enrichment of the band corresponding of a RNA of 17 nts Type A RNA with degradation of 2 nucleotides.

As the NP-exo MOPV presents similar *in vitro* behavior to other Arenavirus NP-exo, we conclude that the ExoN activity *per se* is not responsible for immune suppression, and that the latter is mediated by elements embedded in the domain itself. It was thus of interest to better characterize the substrate specificity of the NP-exo in order to disclose its role in arenavirus replication.

### NP-exo is able to excise a _ds_RNA 3’-end mismatch

We measured NP-exo MOPV and NP-exo LCMV’s ability to cleave different _ds_RNA substrates. Because the key enzyme in the innate immune response; the protein kinase RNA-activated (PKR); is induced by the presence of _ds_RNA, we made use of a perfectly annealed _ds_RNA, as well as several potential RNA substrates such as those mimicking an erroneous replication product with one, two or three mismatched nucleotides at the 3’-end. To that end, a 40-mer RNA template (RT1) blocked in 3’-end with a phosphate group was annealed to a radiolabeled RNA carrying zero (RL2*) or one (RL3*), two (RL4*) or three (RL5*) non-complementary nucleotides at its 3’-end as shown in Fig 4A. Fig 4B shows that both enzymes are strict _ds_RNA ExoNs, with the interesting specific ability to digest substrates carrying a single 3’-terminal mismatch (RL3*/RT1). Quantification of total product (Fig 4C) shows a comparable hydrolytic activity between the perfectly annealed _ds_RNA and a single 3’-terminal mismatch. The cleavage efficiency, however, drastically drops with the number of unpaired bases at the 3’-end.

**Fig 4.**
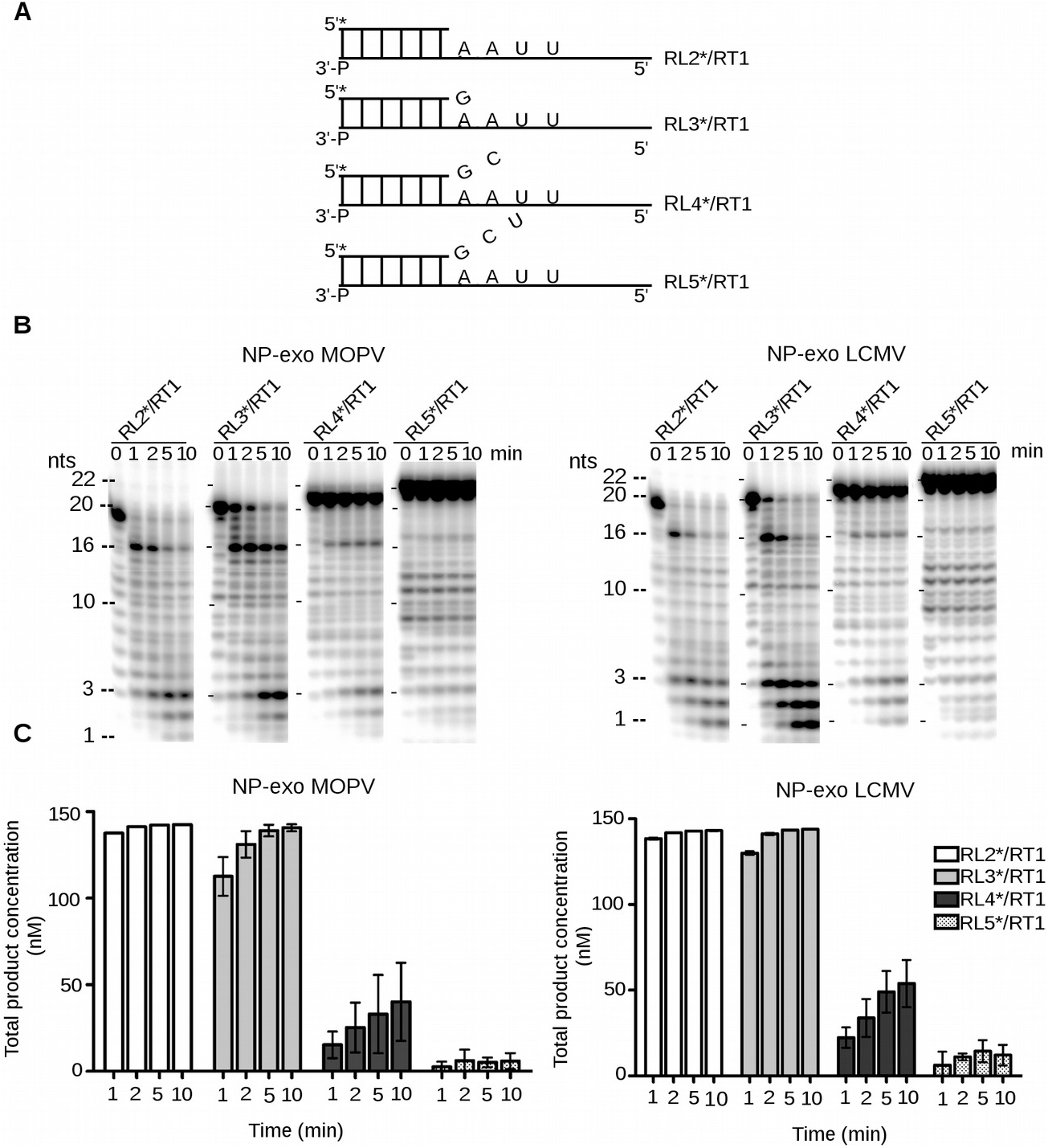
Time course hydrolysis of paired and mismatched 3’-end nucleotide base pair by NP-exo MOPV and NP-exo LCMV. (A) Schematic representation of dsRNA mimicking a replication intermediate. RL2*/RT1, RL3*/RT1, RL4*/RT1 and RL5*/RT1 represent dsRNAs carrying either zero, one, two and three non-complementary nucleotides respectively at their 3’-ends. (B) Equal amounts of the dsRNA substrate (1,25 µM) listed above were incubated in the absence or presence of 0,25 µM of NP-exo MOPV or NP-exo LCMV at 37 °C for 1, 2, 5 and 10 min. 0 is negative control without proteins. A migration size ladder in nucleotides (nts) is presented on the side of the gel. Digestion products were analyzed on 20% denaturing PAGE and visualized by autoradiography. (C) Bar graph showing the total degradation product of substrate at various times. Total degradation products were quantified using phosphoimager FLA-3000 (Fuji) and graphs plotted using Graphpad. PRISM.

### The arenavirus NP-exo domain is structurally and functionally similar to the Coronavirus RNA 3’-mismatch excising ExoN

The overall fold of NP-exo MOPV is homologous to that of the other arenavirus ExoNs [43]. Structural comparison reveals that the structure of NP-exo MOPV is very similar to that of LASV, LCMV and Tacaribe virus (TCRV) structures with overall r.m.s.d of ~1 Å or less, while the residues of the catalytic site are perfectly superimposed. Indeed, the four conserved catalytic residues (D390 E392 D467 H529 D534) from NP-exo MOPV are located at virtually identical positions as those of the other three ExoNs with only minor differences in their orientations. The Zn coordinating residues (E400, C507, H510 and C530) which are highly conserved in arenaviruses are also oriented in an identical manner in all four structures.

A fold similarity search retrieved three ExoN of various origin, namely *Arenaviridae, Coronaviridae*, and a human histone mRNA 3’-ExoNs (S4A Fig). Not only the catalytic core of all these enzymes is conserved (S4B Fig), but also they all possess the ability to remove few unpaired nucleotides in the 3’-to-5’ direction. We analyzed comparatively the NP-exo MOPV structure and the nsp14 SARS-CoV protein (Fig 5) which also possesses a 3’-exoribonuclease activity able to excise 3’-end mismatch on _ds_RNA [47]. The comparison of their topology and of their active site shows that secondary structure elements belonging to the catalytic core are arranged in a similar manner. Our results suggest a common origin of the *Coronaviridae* Nsp14 and *Arenaviridae* NP ExoNs.

**Fig 5.**
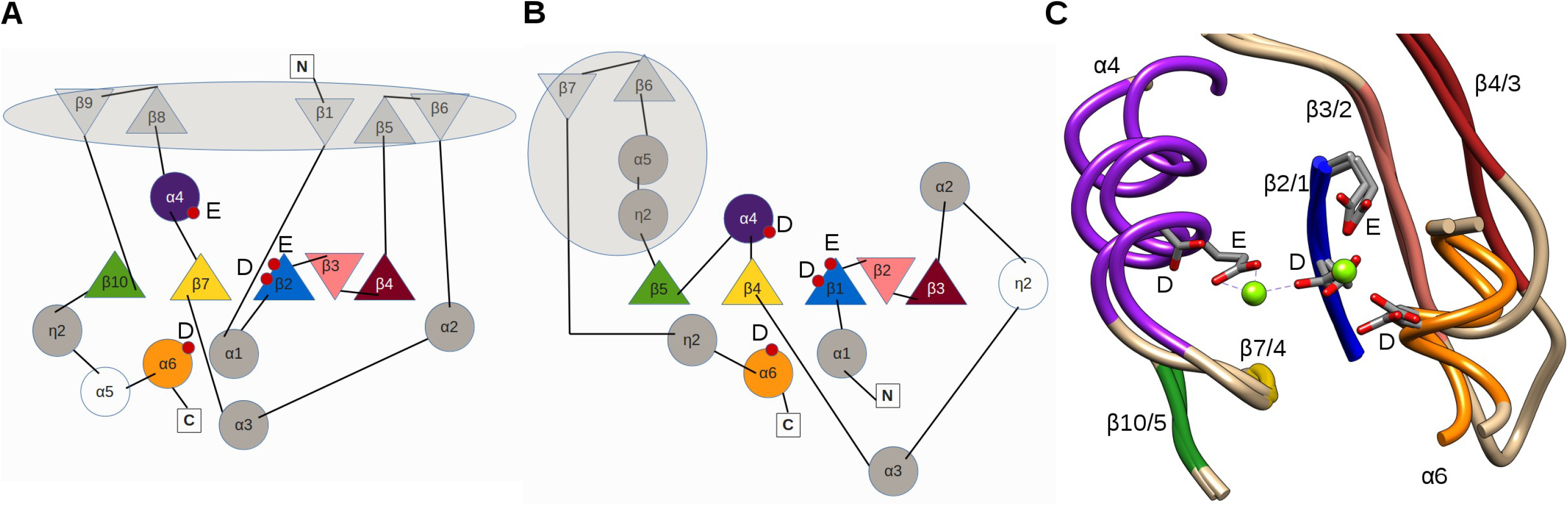
Comparison of the ExoN domains: nsp14 SARScoV and NP-exo MOPV. Topology diagrams of the (A) nsp14 SARScoV and (B) NP-exo MOPV. The 5 β-strands that constitute the central β-sheet are colored blue, pink, brown, yellow and green from the first to the fifth strand respectively. The 4^th^ and 6^th^ α-helices of the DEDD motif are colored purple and orange respectively. The uncolored spheres represent additional non-conserved secondary structures. The location of the catalytic residues are indicated with red spheres. Extra domain insertions for each ExoN are enclosed with in the large gray circles. The N and C terminals are indicated with the uncolored squares. The topology diagrams indicates a similar fold and a conservation of the catalytic core for both ExoNs. (C) Structural alignment of the active sites of nsp14 SARScoV and NP-exo MOPV. Color codes are same as in the topology diagrams except for secondary structures colored cyan (NP-exo MOPV) or sandy brown (nsp14 SARScoV) which are not a part of the catalytic fold. The superposition shows that catalytic residues are located at virtually identical positions.

### The arenavirus NP-exo domain affects viral replication but is not involved in genome stability

To assess the possible role of the NP-exo activity in viral replication and/or genome stability, we passaged iteratively 10 times in Vero E6 cells at a MOI of 0.01 a NP-exo defective virus (NP-exo(-))carrying the D390A/G393A mutants as well as a NP-exo WT recombinant MOPV. We first quantified both the infectious titers (Fig 6A, left axis) and the NP RNA viral loads (Fig 6A, right axis) of the cell culture supernatants. Our results showed that the infectious viral titers of both viruses followed a parallel trend, the NP-exo(-) always presenting a 40 (minimum, passage 4) to 190 (maximum, passage 6) fold decrease in titer compared to the NP-exo WT MOPV. From passages 1 to 5/6, infectious titers continuously decreased (down to a 5 fold for the WT and down a 90 fold for the mutant compared to passage 1) before a rebound from passage 6 to 8/9 to titers similar to those of passage 2 followed by another general decrease at passage 10. The viral loads of NP-exo WT and NP-exo(-) MOPV described the same trends as the infectious viral titers albeit with less pronounced variations. The viral load for the WT virus remained stable along the passages with a maximum 3 fold difference while the NP-exo(-) virus had a maximum 13 fold difference.

**Fig. 6.**
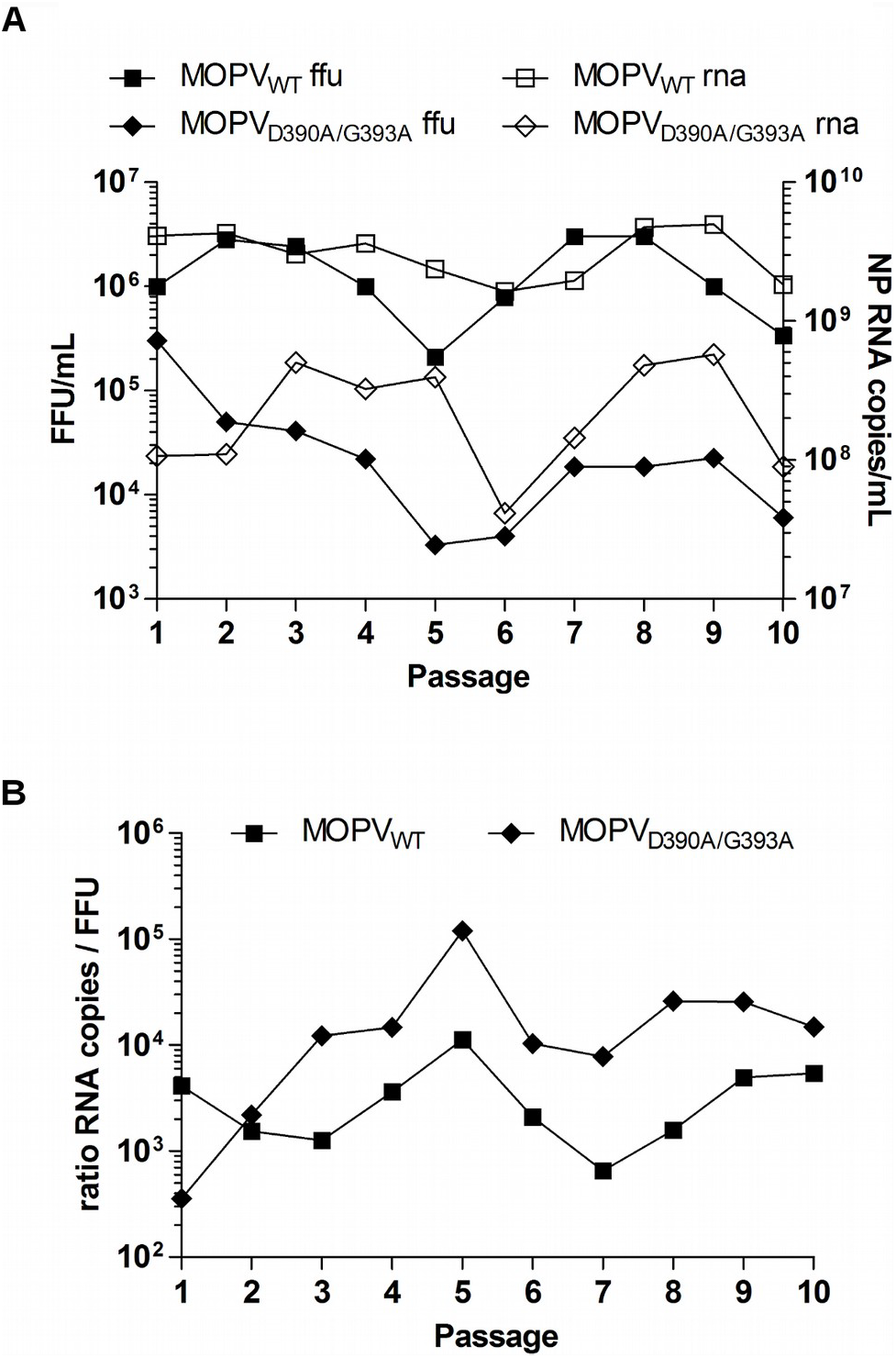
Effect of inactivated ExoN viruses on viral fitness and genome stability. (A) Iterative passages of NP-exo WT and D390A/G393A recombinant MOPV in Vero E6 cells. *De novo* stocks of both viruses, from passage 1 to passage 10, were used to infect cells for 4 days with MOI 0.01. Samples of supernatants were collected for viral infectious titration and NP RNA copy quantification. Results for the left Y axis represent the infectious viral titers (FFU/mL) and results from the right Y axis represent the NP RNA load (NP RNA copies/mL) of the corresponding cell culture supernatants. (B) Ratio calculation for NP RNA copy load over infectious viral titer (RNA copies / FFU) for NP-exo WT and D390A/G393A recombinant MOPV for the ten passages considered in (A).

We also calculated the RNA/FFU ratio for both viruses (Fig 6B). On average, the NP-exo WT and NP-exo mutant viruses respectively presented one infectious particle for 3600 and 23000 NP RNA copies, respectively. Interestingly, the maximum ratio was reached at passage 5 for both viruses with 1 FFU for 11300 copies for the WT NP-exo and 1 FFU for 119600 copies for the NP-exo(-). Therefore, the suppression of the NP-exo activity of MOPV promoted both a reduction of the infectious titer and an increased amount of non-infectious material released from infected cells.

We next investigated the genomic stability of these two viruses at passage 1 and 10 through deep sequencing analysis. We almost reached a complete coverage of the MOPV genomes except for the 5’ and 3’ end and most of the intergenic region of both segments. The tandem repeated and complementary sequences promote strong secondary structures in the intergenic regions and may explain the lack of reads observed for this region for both viruses.

The presence of WT NP sequences detected at passage one for both viruses likely originated from the plasmid expressing the WT NP ORF used for rescuing the virus (data not shown). To make sure our results matched a standard threshold usually observed with the presence of an internal control of sequencing, we considerate a 5% cutoff as a significant read percentage for the effective presence of a mutation (minimum mean coverage of 2636 reads for the L segment of the NP-exo(-) virus and maximum mean coverage of 12610 reads for the S segment of the NP-exo WT virus). As shown in Table 1 & S4, mutations targeting either the ORFs or the UTR/IGR regions are already present in both segments of the NP-exo WT and NP-exo(-) viruses (2.90 and 2.15 mutations/kb respectively) as early as passage one in VeroE6 cells after the virus rescue in BHKT7 cells. The overall mutation rate slightly increases comparatively at passage ten with 3.93 mutations/kb for the NPexo WT virus and 3.36 mutations/kb for the NP-exo(-) virus. Theses results indicate that NP ExoNs does not affect the overall genome stability but affects the viral replication.

**Table 1.**
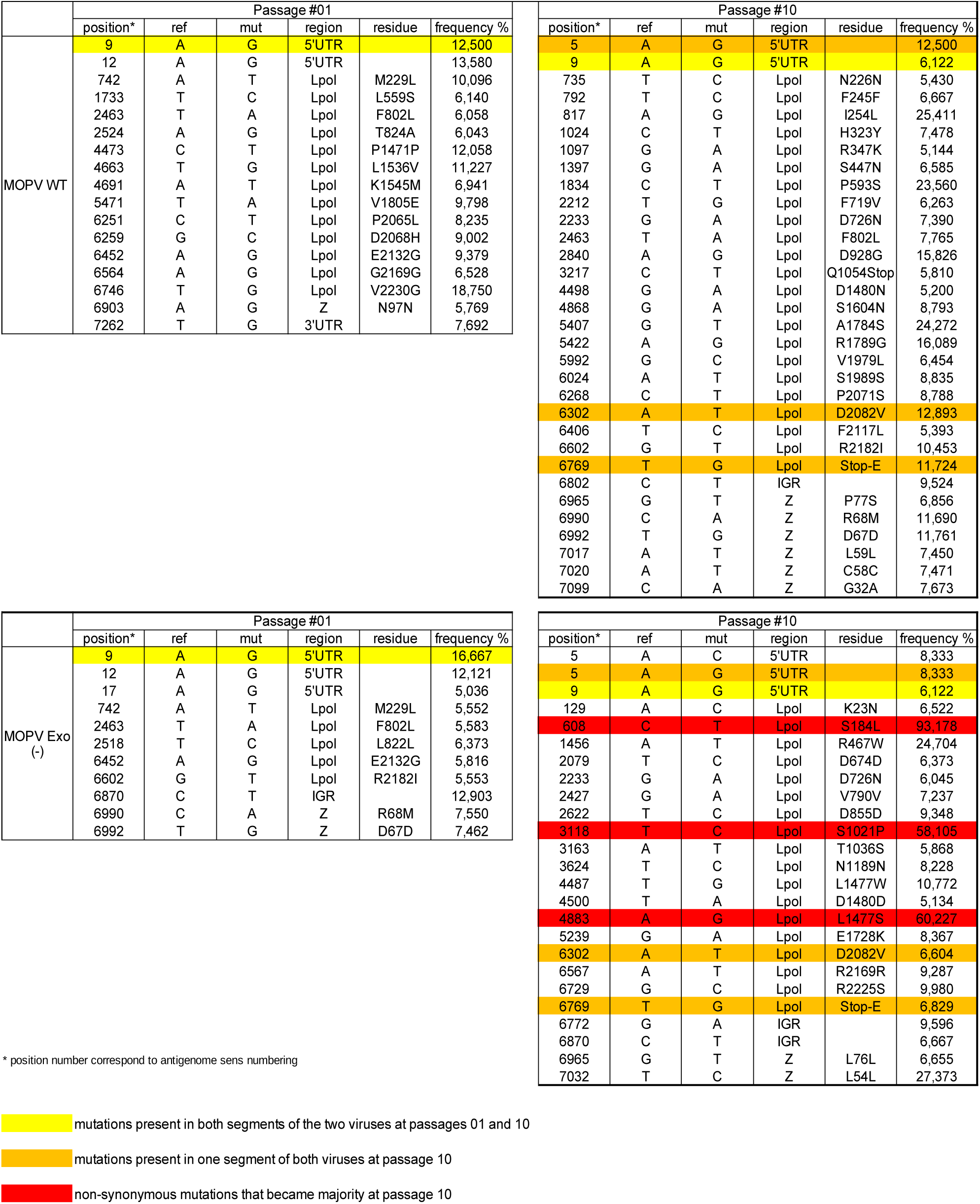
Observed L segment mutations of WT and ExoN(-) viruses at passages #1 and #10.

### The arenavirus NP-exo domain is active on its own genome

We observe that the overall mutation rate is stable between NP-exo WT and NP-exo(-) MOPV, yet we also notice that the frequency of these mutations has changed. We observe at passage 10 a comparable number of mutations along the S segment but a decrease of ~ 22 % on the L segment of the mutant together with an increase of their occurrence frequencies. For both, these mutations appears along the entire L segment at an average frequency of ~ 10.1 % (comprised between 5 to 24 %) for the WT and of ~ 16.6 % (comprised between 5 to 93 %) for the mutant, in particular the substitution of C to T. While for the S segment mutations appear along the entire for the WT and clustered for the mutant at respective frequencies of 14.6 % and 19.5 %.

Among all mutations recorded, only a few were present for either both viruses and/or at the two different passages. Indeed, three mutations in the S segment and one mutation in the L segment are present for both viruses at passages one and ten and likely represent stable quasi-species (Table S4, and Table 1 yellow highlight).

Three mutations in the L segment are commonly found for the two viruses only at passage 10 (Table 1, orange highlight). Interestingly, three non-synonymous mutations in the L-polymerase ORF (S184L, S1021P and L1477S, Table 1, red highlight) became majority for the NP-exo(-) virus at passage 10.

To ascertain the trend observed in our genomic sequencing data, we investigated the natural occurrence of theses mutations in *Mammarenavirus* (MAMV) using bioinformatics. The pre-supposed being that if these mutations appear randomly they should be significantly (> 5 %) represented in the general population of MAMV and LASV.

From the three subsets of sequences generated; *i.e* MAMV, LASV and MOPV; we observed that the three mutations S184L, S1021P and L1477S appeared in conserved regions in all three subsets, that none of the three mutants were reported in MOPV subset and finally that the three amino acid are subject to diverse selective pressure (Fig 7). Indeed, serine at the position 184 represents the majority of the observed amino acid variants while the specific mutation S184L represents only 1 % of the total observed sequences in the LASV or MAMV subsets. This observation indicates that the mutation is viable but most likely costly to be maintained by the virus. On the other hand, at position 1021, the serine observed in MOPV subset does not represent the majority of the observed amino acid in the other subsets, the proline is by far the most frequent amino acids found (Fig 7). In LASV, the serine subset represents only 3% of the observed amino acids at this position while 10% in the MAMV subset (Fig 7). This observation indicates that this position in MOPV is an oddity compared to the others. It seems that the natural tendency in the L protein is to have a proline rather than a serine at this position. The fact that we observe that particular reversion in the NP-Exo(-) could indicate that for L MOPV there is a constraint at this particular position. Finally, at position 1477, the amino acid found at this position for the three subsets is a leucine (Fig 7), indicating that, that particular position is under a high selective pressure. The mutation L1477S was never observed in any subsets, this mutation can be interpreted as unlikely and therefore considered as direct consequence of the NP-Exo(-).

**Fig. 7.**
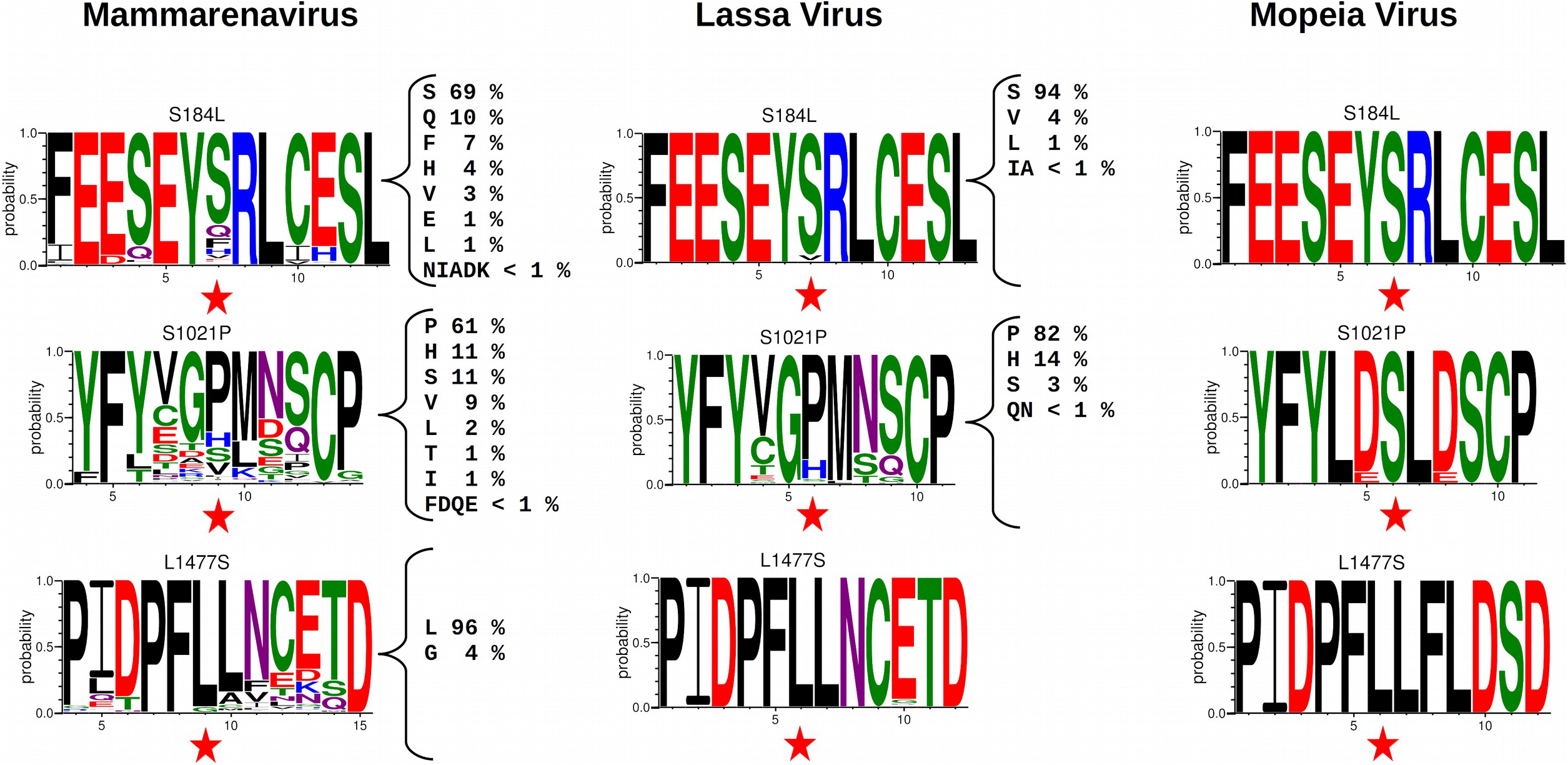
Representation of statistical occurrence of an amino acid at specific position. Represented the WebLogo of the corresponding MOPV amino acids S184, S1021 L1477 in all Mammarenavirus, LASSV and MOPV sub set of sequences. The position of the residue of interest is indicated with a red star. Size of the residue is proportional to it probability of occurrence (0 – 1). For clarity, on the side of the WebLogo is the detailed the statistical occurrence of all amino acids found in the deposited sequences.

Our results show that compared to the WT virus, the abrogation of the NP-exo activity did not increase the mutation rate found in the MOPV genomic sequences present in the cell culture supernatant, but the emergence of the three mutants of rare occurrence are the direct consequence of the NP-Exo(-), which have relaxed the control over certain position implying a direct effect of the MOPV exonuclease on its own genomic RNA.

## Discussion

The paradigm of *Arenaviridae* NP ExoN states that it is involved in innate immunity suppression [37,60–62], through degradation of _ds_RNAs which would otherwise stimulate the innate immunity response. Several reports have demonstrated that NP is responsible for the degradation of these _ds_RNAs using the 3’-5’ ExoN located at its C-terminus [25,32,35,63]. This ExoN comprises a DEDDh catalytic motif that is completely conserved across the *Arenaviridae* [35] implying this activity may be a general feature of arenavirus NPs. The ExoN domain is conserved within the family regardless of both the virus pathogenic potential and its ability to suppress efficiently type I IFN, as previously reported for TCRV and MOPV [42,60,64]. MOPV is the closest counterpart of LASV and presents a 73% NP sequence identity with LASV. During LASV infections, the virus targets mainly macrophages (MP) and dendritic (DC) cells [65], and infections are characterized by high viremia and generalized immune suppression supposedly due to innate immune inhibition by the ExoN domain. Both MOPV and LASV induces strong type I IFN responses in MP and moderately in DC, but contrarily to LASV which abrogates this response, MOPV does not [42,64].

Our functional study demonstrates that the ExoN structure, substrate specificity, and mechanism is indeed conserved across the family. It is also clear from the structure-ion analysis [43], that toying with the catalytic ion leads to slight structural changes which impact dramatically the activity. Although NP-exo MOPV, NP-exo LCMV have similar cleavage patterns, mutation analysis of the DEDDh motif reveals that the exact residues critical for 3’-5’ ExoN vary between both domains. For NP-exo MOPV, D390A, E392A and D534A (D389A, E391A and D533A LASV equivalents) completely abolishes 3’-5’ ExoN activity consistent with results from *in vitro* studies on LASV [35], while D467A and H529A (D466A and H529A LASV equivalents) retains some residual activity (see below). For NP-exo LCMV, a previous study by Martínez-Sobrido and collaborators [61], correlated innate immunity suppression to ExoN mutants, postulating a direct involvement of the ExoN activity. We observed that mutant D382A completely loses ExoN activity consistently with results from reverse genetic studies [61]. A noticeable difference concern the mutant E384A, that was shown to have no effect and be dispensable for ExoN activity [61], is rather shown critical in our *in vitro* study and consistent with the structural data as E384 (equivalent to MOPV E392) is involved in binding one of the catalytic ion (S4 Fig). Under our conditions, D459A and H517A still retains their ability to cleave two nucleotides meanwhile D522A shows a significant activity leading to the removal of two to three nucleotides. Theses latter three mutants were not reported before but the analysis of the structure of NP-exo MOPV confirms that major features such as fold, and the two ion binding sites (catalytic and structural) are conserved within the *Arenaviridae* [25,32,35,36,63]. Residues D390, E392, D534 of NP-exo MOPV directly coordinate the catalytic ion. Mutation of these residues logically alter ion binding and thus leads to complete loss of catalytic activity. The residual activity observed for D522A of NP-exo LCMV is rather difficult to explain as the two structures present no clear differences in the ion binding mode. The only noticeable difference is about the hydrophobic environment that may compensate the faulty metal-binding site in the case of NP-exo LCMV. The residual activity observed for D467 (D459 of LCMV) is rather difficult to explain as the two structures present no clear differences, yet that residue is very likely involved in the cleavage mechanism. Indeed, the kinetic experiments comparing the WT NP-exo MOPV and NP-exo LCMV to corresponding mutants show that the first event of the hydrolysis is comparable between NP-exo MOPV and NP-exo LCMV, while mutation of the aspartate reduces drastically the hydrolysis kinetics for NP-exo MOPV but only moderately for NP-exo LCMV (S3 Fig). These differences suggest that the aspartate (respectively D467, D459) is involved in the structural set-up of the active site for positioning the ion responsible for the nucleophilic attack. The general mechanism for RNA hydrolysis is a two metal ion mechanism described by Steitz and Steitz [66]. It involves metal A and B positioned ~4 Å apart each other and across the target phosphodiester bond. Metal ion A facilitates the formation of the attacking nucleophile. This is followed by the formation of a penta-covalent intermediate which is stabilized by both metal ions. Metal ion B then eases the exit of the leaving group. With the help of the LASV structures of Jiang and collaborator [36], we made an attempt to reconstitute a model of the general mechanism for RNA hydrolysis (S5A Fig). In NP-exo MOPV structure (and all the others) only the metal B is visible. The site receiving the other catalytic metal A is partly created with interaction of the RNA and residues D390 and D467 (S5B Fig). The interaction between the ion in position A and D467 is mediated through a water molecule. Therefore, this might explain why this mutant retained a partial residual activity.

As it was shown for TCRV, the ExoN domain of MOPV *in vitro* is endowed with full ExoN activity and obeys to the same structural and energetic constraints as those of other *Arenaviridae* ExoNs [36,43]. Therefore MOPV ExoN activity alone is not *per se* responsible for the differences in innate immunity suppression between MOPV and LASV. Rather, the presence of the ExoN activity may serve other purposes in the viral life cycle, which might be connected directly or indirectly to innate immunity.

In particular, previous studies on LCMV and PICV suggested that altering the NP ExoN also impacts replication, irrespective to the IFN status of the host cell [61,67]. The structural relatedness with the Coronavirus ExoN and its implication in viral replication prompted us to investigate to which extend the arenavirus ExoN domain is able to excise unpaired nucleotides. Our enzyme activity assays demonstrate that NP-exo MOPV and NP-exo LCMV can efficiently and selectively cleave a _ds_RNA mimicking an erroneous replication product carrying one 3’-mismatched nucleotide (Fig 4). Our analysis confirms that despite additional inserted structural elements, the two domains belong to the same Ribonuclease H-like superfamily. In the case of *Coronaviridae,* several studies have pointed to a main role of the ExoN domain of nsp14 in RNA proofreading [47–50] to maintain genome stability. Structural comparison between ExoN domain of nsp14 and MOPV shows conservation of active site and main fold (Fig 5) suggesting that they have a distant but common origin. Recent work by Becares and colleagues have shown that nsp14 of Coronavirus is also involved in innate immunity modulation [68]. Therefore, these data show that at least in *Coronaviridae* the 3’-5’ ExoN activity is not exclusively assigned to a specific role but is involved in different aspect of the viral life cycle. Our study present clear evidence that much like the coronavirus nsp14 [47], the *Arenaviridae* NP ExoN excises a 3’-end mismatch _ds_RNA *in vitro*, and based on previous report that this activity is directly connected to RNA replication [61,67].

From our data, the MOPV polymerase exhibits an average error rate estimated around 3 mutations / kb. This means that the polymerase was able to incorporate a mismatch and then elongate it. The error rate does not change between the WT and the mutant, which is consistent with the fact that we did not altered the polymerase. For both, these mutations appears along the entire L segment at an average frequency of ~10.1 % (comprised between 5 to 24 %) for the WT and of ~16.6 % (comprised between 5 to 93 %) for the mutant. The fact that unlikely mutants have become prevalent, as observed at passage 10, reflect that the control over certain type of mutation have been abolished, thus implicating that ExoN is active on its own genomic RNA. In this study, it is not the particular set of mutations that is relevant but rather that a set of unlikely mutations have emerged. The bias inferred by impairment of the ExoN, together with the biochemistry presented here is consistent with the idea that ExoN is involved in a mismatch excision system.

Although the presence of mismatch excision system is logically associated to very large genomes (~30 kb) in *Coronaviridae*, the presence of such activity, and potentially such RNA repair system, in *Arenaviridae* of intermediate genome size (~11 kb) remains puzzling. Our results shows a clear diminution of the viral titer for viruses depleted of ExoN activity, but no clear evidence of a drastic increase of mutation in the genome that would lead to catastrophic event. Then what is happening in these mutated viruses? One tentative explanation could be the ExoN is involved in: *i*) checking and maintaining the sequence integrity of the conserved genomic region at its extremities, and/or *ii*) the structural integrity of the Intergenic Region (IGR). Indeed, both regions have been previously reported as being critical for viral fitness: *i*) The conserved region (19 nucleotides) exhibit high degree of sequence conservation at the 3’-termini and is complementary to the 5’ end of the genome (for review [26]). This sequence serves as a selective docking platform for the polymerase [69] for which 3’-end binds with high affinity and in a sequence specific manner [70]. *ii*) Similarly, alteration of the IGR structure leads to reduce efficient transcription termination and viral assembly [71,72]. In such hypothesis, the impairment of the NP-exo activity leads to a scenario in which the polymerase is able to incorporate a mismatch but is unable to elongate it, leading to a decrease of suitable genomic material to package, therefore without the ExoN control the number of functional RNP would be reduced and consistent with the loss of viral fitness observed here and else [61,67]. Therefore we propose that the Arenavirus ExoN is involved in a “limited proof-reading” mechanism driven by structural constraints rather than genomic stability. Another observation that concurs to the “limited proof-reading” mechanism is the difference of Ribavirin efficiency on *Arenaviridae and Coronaviridae.* Ribavirin is the only drug so far administered on large scale and having demonstrated a decrease of mortality rates up to 5%, if administered within the first 6 days of arenaviral illness [73]. On the other hand, Ribavirin is ineffective against coronaviruses [74,75], as nsp14 ExoN domain excises the nucleotide analogue [51]. It is likely, that for *Arenaviridae,* the ExoN activity involved in a “limited proof-reading” mechanism remains as a trace of its past common ancestor with *Coronaviridae*. The critical problem of genomic stability being solved, either by the conservation of the “original” function of the ExoN for *Coronaviridae*, or by genome segmentation for *Arenaviridae*.

As a conclusion we have shown that the MOPV ExoN is fully functional, behaving like other Arenaviral ExoN on _ds_RNA. We have demonstrated that Arenaviral ExoN are able to excise an RNA mismatch, and that is active on its own genomic RNA like its counterpart in *Coronaviridae*. Under the conditions used here, abrogation of ExoN activity does not impact genomic stability significantly. Our results suggest that the *Arenaviridae* RNA ExoN, like that of *Coronaviridae*, is at a crossroad between replication efficiency and innate immunity evasion in the infected cell.

## Supporting information

Supplemental files ant Tables

## Acknowledgments

This work was supported by ANR grants ArenaBunya-L (ANR-11-BSV8-0019), the Centre National de la Recherche Scientifique (CNRS), the Fondation Infection Mediterrannée, and the French Infrastructure for Integrated Structural Biology (FRISBI) ANR-10-INSB-05-01. The authors also thank Dr. Barbara Selisko for her help and useful discussion and Julie Lichière for her contribution in the experiments.

## Supporting information

**S1 Fig. Modular and organization of *Arenavidae’* NP**. Schematic of the two domains organization of *Arenaviridae*’ NP, with its corresponding domains structures. N-terminal domain corresponding to the nucleoprotein domain (PDB:3T5N) and C-terminal domain corresponding to the ExoN domain (PDB:3Q7C) of LASV. Each domain is represented in ribbon and colored following rainbow nomenclature from blue (N-terminus) to red (C-terminus). Flexible linker is represented as green line.

**S2 Fig. Secondary structures adopted by RNA substrates used in the study**. (A) These structures were predicted using Mfold server (http://unafold.rna.albany.edu/?q=mfoldand). The minimum free energy of each structure is indicated below it. RNA HP4 is a stable RNA hairpin. RNA LE19 presents 3 types of secondary structures. Type A: two double stranded nucleotides, type B: long hanging 5’ and short hanging 3’ extremities, type C: long 5’ 3’ Hanging extremities. RNA A30 long single stranded RNA without secondary structure. (B) Expected pattern of digestion based on a dsRNA 3’5’ ExoN activity for each type of tested RNA.

**S3 Fig. Kinetics of HP4 cleavage by NP-exo MOPV WT, its Mutant D467A, NP-exo LCMV WT or the corresponding mutant D459A.** HP4 was incubated with equal concentration of NP-exo MOPV WT, D467A, NP-exo LCMV WT or D459A for 0, 2, 4, 6, 8, 10 and 15 mins. Reactions products were separated on 20 % denaturing PAGE and products revealed by autoradiography. RNA cleavage was then quantified from this data and plotted as the percentage of product formed with time.

**S4 Fig. Structural alignment of 3’-5’ ExoN**: (A) Ribbon representation of the superposition of NP-exo Mopeia Virus (cyan pdb:5LRP), nsp14 SARScoV Exo (beige pdb:5C8S), NP-exo Lassa Virus (orange pdb:4FVU), histone mRNA stem-loop by 3’-ExoN Homo Sapiens (green pdb 1ZBH). All structures were retrieved by PDBeFold (http://www.ebi.ac.uk/msd-srv/ssm), all similar sequences were discarded. Overall fold is similar; in caption a zoom of the active site shows a quasi perfect superimposition of the DEDD motif. Green spheres are in the zoom represent Mg ions. (B) Weblogo of the active site derived from the structural alignment. Structure and sequence comparison lead to conclude to a common origin of the ExoNs.

**S5 Fig. A model of the mechanism of calcium inhibition of 3’-5’ ExoN activity.** (A) The dsRNA from LASV NP-C structure (PBD code:4GV9) modeled into the MOPV Nexo-Mg structure. MOPV-Mg represented as cartoon (Helices in orange, β-strands in green and loops in cyan) and the dsRNA is shown as a stick model. The green balls represent magnesium atom. (B) An enlarged view of the active indicating the model positions of ions during cleavage mechanism. Calcium substitution of the magnesium ion mediates inhibition of 3’-5’ ExoN activity as a result of its atomic radius, binding flexibility and poor activation of water. Ca atom is represented in grey.

**S1 Table. Primer sequences used for mutagenesis of the LCMV and MOPV.**

**S2 Table. Oligomers names and sequences.**

**S3 Table. Comparison of remaining sequence length function of time.**

**S4 Table. Observed S segment mutations of WT and ExoN(-) viruses at passages #1 and #10.**

