## Supplemental files ant Tables for "*Arenaviridae* exoribonuclease presents genomic RNA edition capacity"

### FigS1

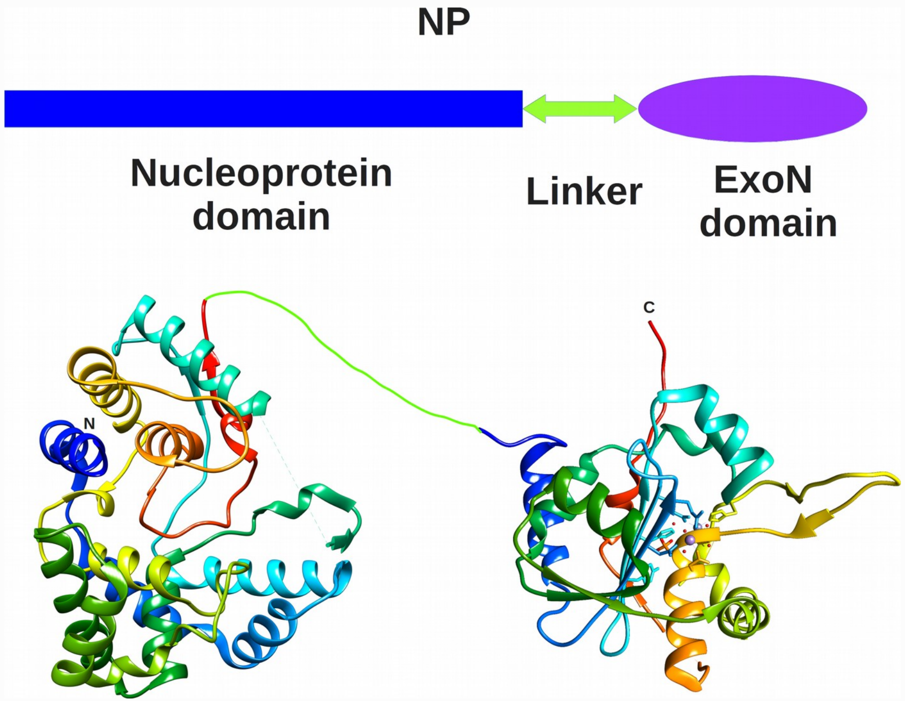

### FigS2

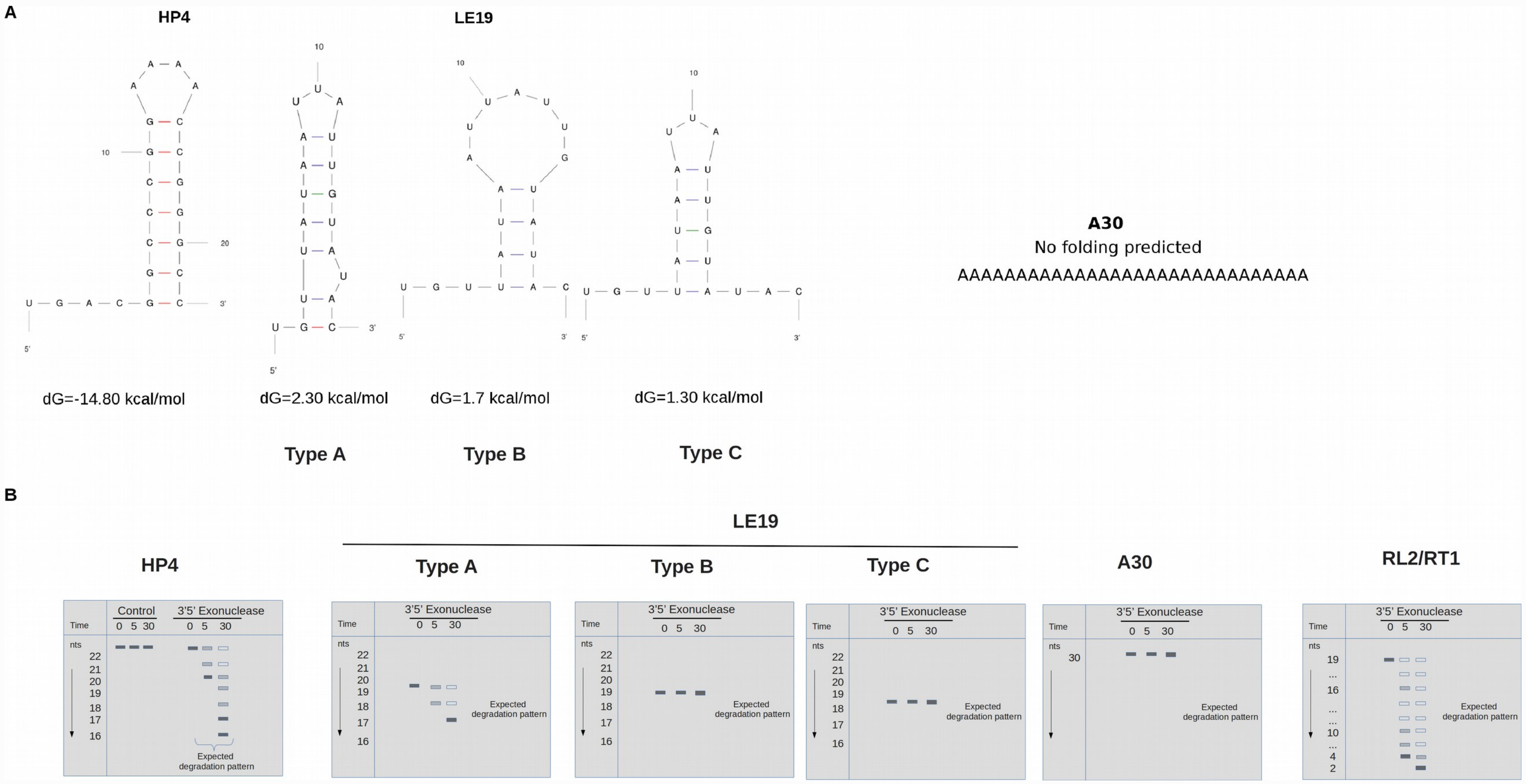
A
HP4
LE19
dG=-14.80 kcal/mol
dG=2.30 kcal/mol
dG=1.7 kcal/mol
dG=1.30 kcal/mol
Type A
Type B
Type C
A30
No folding predicted
AAAAAAAAAAAAAAAAAAAAAAAAAAAAAA
B
LE19
HP4
Type A
Type B
Type C
A30
RL2/RT1
Control
3'5' Exonuclease
Time
0 5 30
nts
Expected degradation pattern

### FigS3

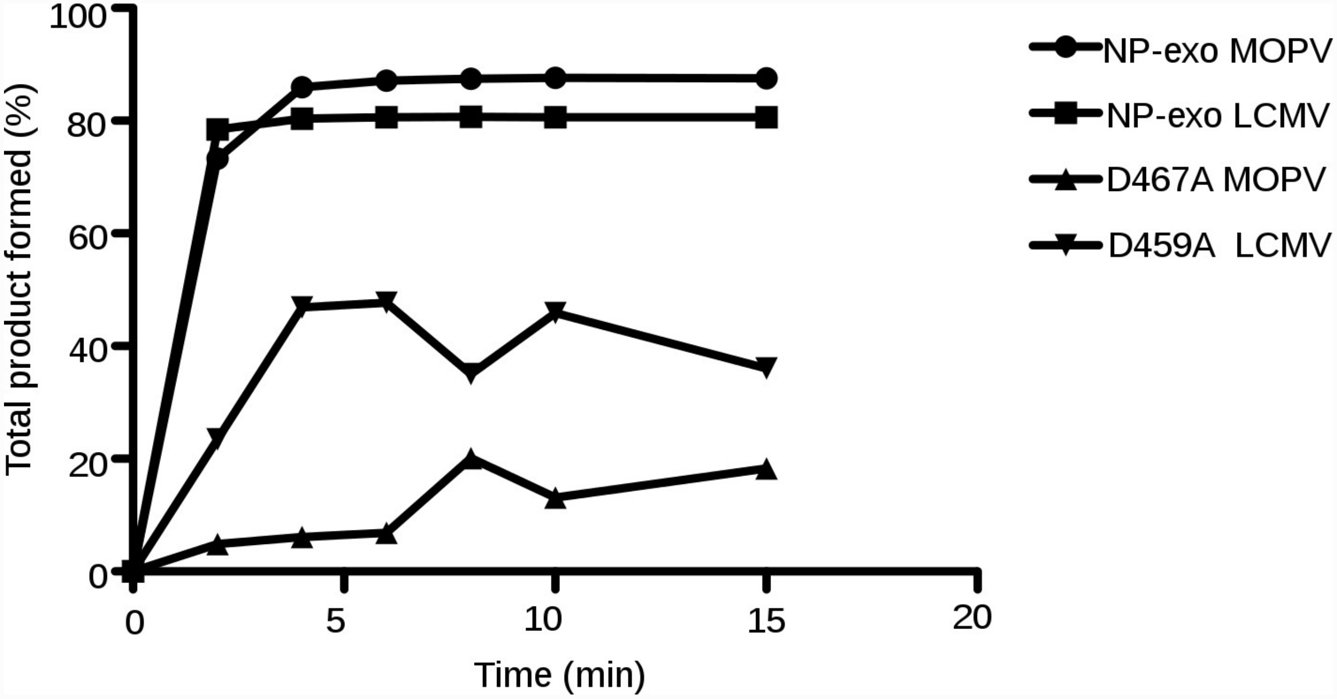

100
80
60
40
20
0
Total product formed (%)
0
5
10
15
20
Time (min)
NP-exo MOPV
NP-exo LCMV
D467A MOPV
D459A LCMV

### FigS4

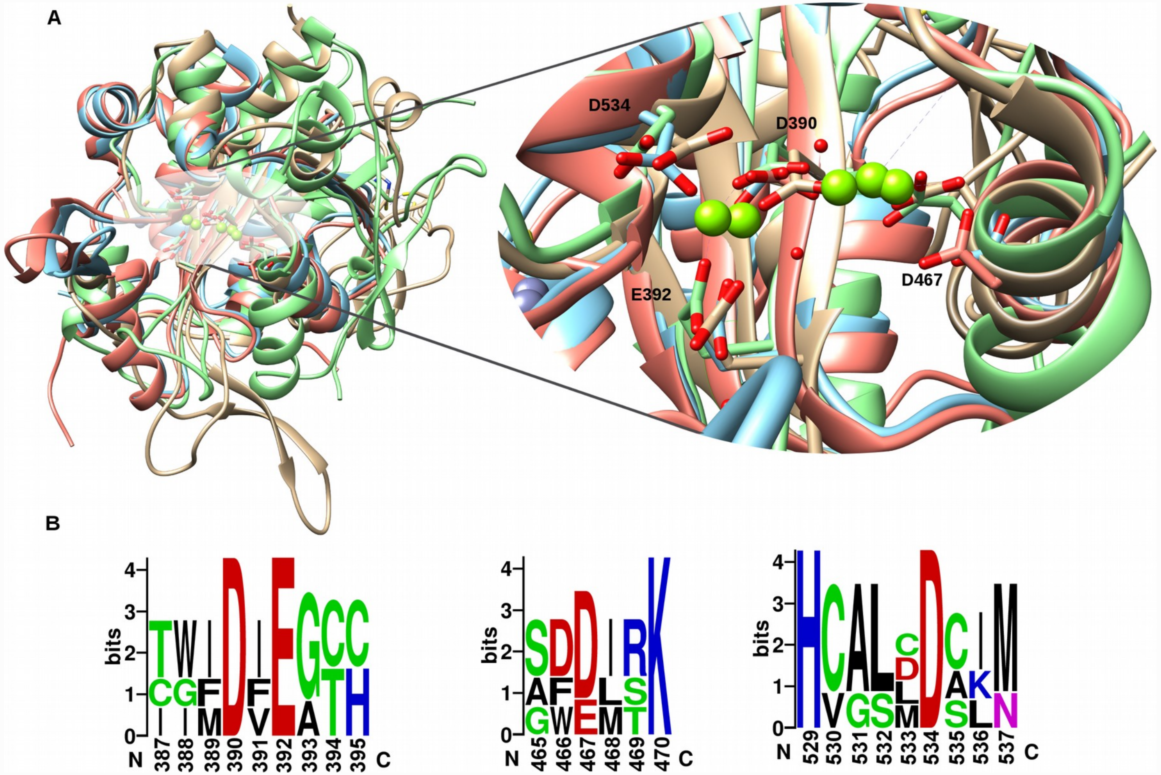
A
D534
D390
E392
D467
B
bits
N
387 388 389 390 391 392 393 394 395
C
465 466 467 468 469 470
529 530 531 532 533 534 535 536 537

### FigS5

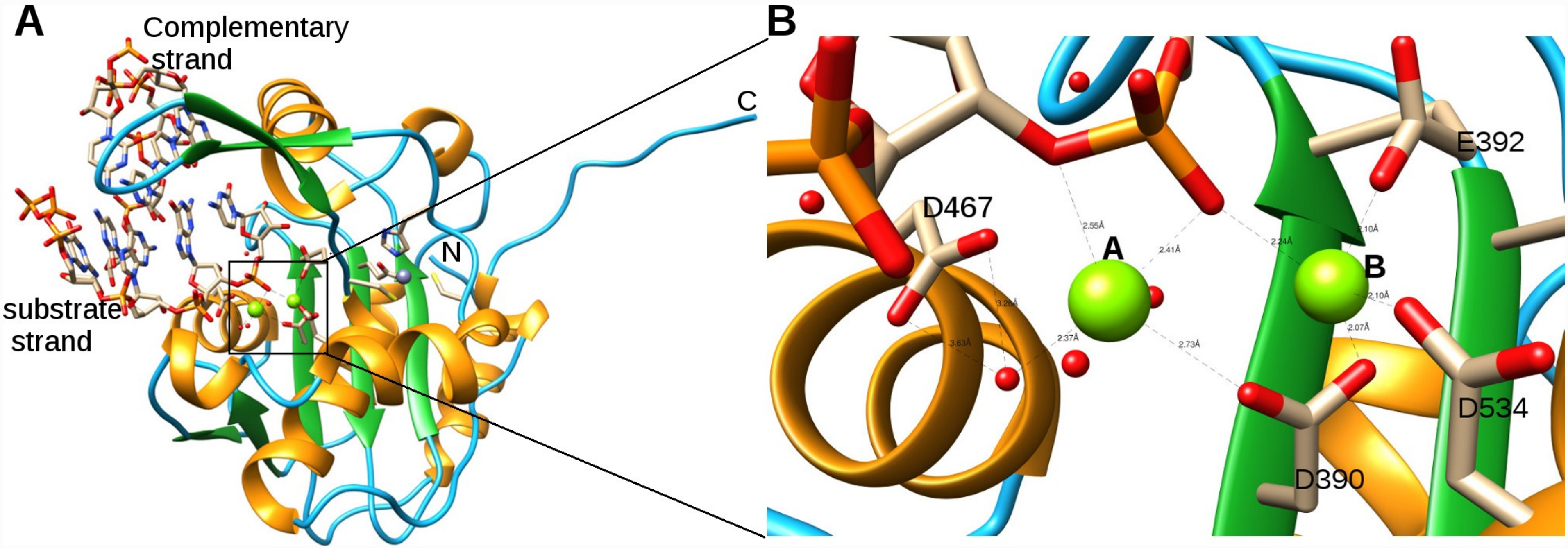
A
Complementary strand
substrate strand
C
N
B
D467
E392
A
B
D534
D390

| LCMV | Primer sequences |
|---|---|
| D382A_R | atcattaaatctaccctcaatgGcaatccatgtaggagcgttg |
| D382A_F | caacgctcctacatggattgCcattgagggtagatttaatgat |
| D459A_R | gagtccagaagctttctgatTGCatcggagccttgacagctta |
| D459A_F | taagctgtcaaggctccgatGCAatcagaaagcttctggactc |
| D517A_R | tgcagtccatgagtgcacagGCcggggtgatctctttcttcttt |
| D517A_F | aaagaagaaagagatcaccccgGCctgtgcactcatggactgca |

| MOPV | Primer sequences |
|---|---|
| D390A_AS | cttcaggtctcccttctatgGcaatccatgtcttagcattt |
| D390A_S | aaatgctaagacatggattgCcatagaagggagacctgaag |
| E392A_AS | ggtcttcaggtctccctGctatgtcaatccatgtc |
| E392A_S | gacatggattgacatagCagggagacctgaagacc |
| D467A_AS | tcaagaagctttctgatgGcatcagcaccttgacacg |
| D467A_S | CgtgtcaaggtgctgatgCcatcagaaagcttcttga |
| D534A_AS | tgcttcaaacatgagacaaGccagcaatgcacagtgtg |
| D534A_S | cacactgtgcattgctggCttgtctcatgtttgaagca |
| H529A_AS | caatccagcaatgcacagGCtggtgtcacttcttcttt |
| H529A_S | aaagaagaagtgacaccaGCctgtgcattgctggattg |

***S1 Table :* Primer sequences used for mutagenesis of the LCMV and MOPV.**

| Oligomer name | Sequence |
| --- | --- |
| HP4 | 5'-UGA CGG CCC GGA AAA CCG GGC C-3' |
| LE19 | 5'-UGU UAU AAU UAU UGU AUA C-3' |
| A30 | 5'-AAA AAA AAA AAA AAA AAA AAA AAA AAA AAA-3' |
| RT1 | 5'-CUAUCCCCAUGUGAUUUUAAUAGCUUCUUAGGAGAAUGAC-3' |
| RL2 | 5'-GUCAUUCUCCUAAGAAGCUA-3' |
| RL3 | 5'-GUCAUUCUCCUAAGAAGCUAG-3' |
| RL4 | 5'-GUCAUUCUCCUAAGAAGCUAGC-3' |
| RL5 | 5'-GUCAUUCUCCUAAGAAGCUAGCU-3' |

***S2 Table : Oligomers names and sequences.***

| **WT MOPV** | Substrate length (nts) / t (min) | | | |
|---|---|---|---|---|
| Ions / t (min) | 0 | 0.1 | 5 | 30 |
| $Mg^{2+}$ | 22 | 21 | 18 | 18 |
| $Mn^{2+}$ | 22 | 20 | 13 | 13 |
| $Zn^{2+}$ | 22 | 22 | 22 | 22 |
| $Ca^{2+}$ | 22 | 22 | 22 | 22 |
| EDTA | 22 | 22 | 22 | 22 |
| **WT LCMV** | Substrate length (nts) / t (min) | | | |
| Ions / t (min) | 0 | 0.1 | 5 | 30 |
| $Mg^{2+}$ | 22 | 20 | 17 | 17 |
| $Mn^{2+}$ | 22 | 20 | 15/14 | 14/13 |
| $Zn^{2+}$ | 22 | 22 | 22 | 22 |
| $Ca^{2+}$ | 22 | 22 | 22 | 22 |
| EDTA | 22 | 22 | 22 | 22 |

***S3 Table : Comparison of remaining sequence length function of time.***

| | Passage #01 | | | | | | Passage #10 | | | | | |
|---|---|---|---|---|---|---|---|---|---|---|---|---|
| | position* | ref | mut | region | residue | frequency % | position* | ref | mut | region | residue | frequency % |
| MOPV WT | 48 | C | G | 5'UTR | | 26,0 | 48 | C | G | 5'UTR | | 19,9 |
| | 52 | A | G | 5'UTR | | 16,1 | 52 | A | G | 5'UTR | | 12,8 |
| | 86 | A | G | NP | E5G | 5,6 | 84 | G | A | NP | K5K | 22,1 |
| | 202 | T | A | NP | S45T | 6,5 | 1057 | G | C | NP | A330P | 5,1 |
| | 404 | A | G | NP | K112R | 5,9 | 1732 | T | G | NP | I548S | 11,0 |
| | 481 | A | G | NP | R138G | 6,9 | 1772 | T | G | NP | V568G | 51,0 |
| | 1386 | A | G | NP | L439L | 5,6 | 1777 | C | G | NP | L570V | 5,4 |
| | 1393 | G | A | NP | V442I | 5,2 | 2761 | T | G | GPC | I205I | 5,8 |
| | 1772 | T | G | NP | V568G | 40,9 | 2963 | G | C | GPC | T138R | 5,1 |
| | 1793 | C | A | IGR | | 22,2 | 3380 | G | T | 3'UTR | | 7,7 |
| | 1916 | C | T | IGR | | 9,7 | | | | | | |
| | 2269 | G | C | GPC | Y369Stop | 5,7 | | | | | | |
| | 3324 | C | T | GPC | V18M | 6,7 | | | | | | |
| | 3376 | C | T | 3'UTR | | 10,1 | | | | | | |

| | Passage #01 | | | | | | Passage #10 | | | | | |
|---|---|---|---|---|---|---|---|---|---|---|---|---|
| | position* | ref | mut | region | residue | frequency % | position* | ref | mut | region | residue | frequency % |
| MOPV Exo (-) | 48 | C | G | 5'UTR | | 6,9 | 33 | C | T | 5'UTR | | 8,0 |
| | 52 | A | G | 5'UTR | | 7,8 | 48 | C | G | 5'UTR | | 21,4 |
| | 321 | A | T | NP | Q84H | 6,3 | 52 | A | G | 5'UTR | | 12,8 |
| | 356 | C | T | NP | S96F | 8,0 | 1632 | T | C | NP | G521G | 50,6 |
| | 682 | T | A | NP | L205M | 5,8 | 1697 | T | C | NP | I543T | 48,8 |
| | 1386 | A | G | NP | L439L | 6,5 | 1732 | T | G | NP | I548S | 11,2 |
| | 1393 | G | A | NP | V442I | 6,0 | 1772 | T | G | NP | V568G | 26,9 |
| | 1652 | C | G | NP | P528R | 7,3 | 1793 | C | A | IGR | | 5,2 |
| | 1772 | T | G | NP | V568G | 48,1 | 1804 | A | C | IGR | | 11,1 |
| | 1793 | C | A | IGR | | 10,5 | 1996 | A | G | GPC | P460P | 11,4 |
| | 1899 | C | G | IGR | | 12,9 | 3224 | A | G | GPC | F51S | 7,0 |
| | 3224 | A | G | GPC | V18M | 5,2 | | | | | | |

(yellow) mutations present in both segments of the two viruses at passages 01 and 10

* position number correspond to antigenome sens numbering

***S4 Table : O*bserved mutation between WT and Mutant Exo (-) of segment S at passages #1 and #10.**
